# JINXED: Just in time crystallization for easy structure determination of biological macromolecules

**DOI:** 10.1101/2022.10.26.513656

**Authors:** Alessandra Henkel, Marina Galchenkova, Julia Maracke, Oleksandr Yefanov, Johanna Hakanpää, Jeroen R. Mesters, Henry N. Chapman, Dominik Oberthür

**Affiliations:** Center for Free-Electron Laser Science CFEL, Deutsches Elektronen-Synchrotron DESY, Notkestr. 85, 22607 Hamburg, Germany; Deutsches Elektronen-Synchrotron DESY, Notkestr. 85, 22607 Hamburg, Germany; Institut für Biochemie, Universität zu Lübeck, Ratzeburger Allee 160, 23562 Lübeck, Germany; Department of Physics, Universität Hamburg, Luruper Chaussee 149, 22761, Hamburg, Germany; The Hamburg Center for Ultrafast Imaging, Universität Hamburg, Luruper Chaussee 149, 22761, Hamburg, Germany

## Abstract

Macromolecular crystallography is a well-established method in the field of structure biology and has led to the majority of known protein structures to date. After focusing on static structures, the method is now developing towards the investigation of protein dynamics through time-resolved methods. These experiments often require multiple handling steps of the sensitive protein crystals, e.g. for ligand soaking and cryo-protection. These handling steps can cause significant crystal damage, causing a decrease in data quality. Furthermore, in time-resolved experiments based on serial crystallography that use micron-sized crystals for short diffusion times of ligands, certain crystal morphologies with small solvent channels can prevent sufficient ligand diffusion. Described here is a method combining protein crystallization and data collection in a novel one-step-process. Corresponding experiments were successfully performed as a proof-of-principle using hen egg white lysozyme and crystallization times of only a few seconds. This method called JINXED (<u>J</u>ust <u>in</u> time <u>c</u>rystallization for <u>e</u>asy structure <u>d</u>etermination) promises to result in high-quality data due the avoidance of crystal handling and has the potential to enable time-resolved experiments with crystals containing small solvent channels by adding potential ligands to the crystallization buffer, simulating traditional co-crystallization approaches.

## Introduction

Protein crystallization was first described more than 150 years ago (Giegé, 2013; McPherson, 1991; McPherson and Gavira, 2014) and has enabled the structure determination of biological macromolecules at an atomic level (Dickerson, 2005). Notwithstanding the recent success of single molecule cryoEM, crystallography is still the method of choice in many areas of structural biology and structure-based drug discovery. The crystallization of biological macromolecules is a tedious process – owing to the complex structure, dynamic behaviour, and complex surface charge distribution of said molecules. Even after many decades of research it is not completely understood. Once crystals of good quality are obtained, the standard single-crystal rotational data collection at cryogenic temperatures requires a lot of crystal handling from fishing, ligand soaking, cryo-protecting to flash-cooling. These various handling steps often damage the sensitive protein crystal, reducing its quality and diffraction capabilities and hence limit the chances of collecting high resolution diffraction data (Dobrianov et al., 1999). Thus, a lot of research focused on mitigating these effects, mainly through the development of *in-situ* crystallization approaches, aided in part by the coming of age of microfluidic methods from the middle of the 1990s onwards (Hansen and Quake, 2003; Heymann et al., 2014; Perry et al., 2013; De Wijn et al., 2019; Yadav et al., 2005). With serial crystallography (Barends et al., 2022; Boutet et al., 2012; Chapman et al., 2011) and subsequent room-temperature data collection coming into play, not only has the required crystal size decreased to micron and sub-micron dimensions but also required handling steps were drastically reduced since fishing, cryo-protection, and flash cooling have become unnecessary. Furthermore, structural biology no longer exclusively focuses on static structures (Martin-Garcia, 2021) but – as biology naturally implies dynamical interactions between molecules – aims to reveal protein dynamics, especially when it comes to protein-ligand interactions. Serial crystallography offers the possibility to study these interactions at an atomic level in time-resolved experiments (Brändén and Neutze, 2021; Mehrabi et al., 2019; Pande et al., 2016; Stagno et al., 2017; Tenboer et al., 2014), e.g. by mixing the crystalline slurry with substrate containing solution before probing the mixture with X-rays after a defined time delay (Beyerlein et al., 2017). However, chemical mixing is limited by crystal and solvent-channel sizes, accessibility of the binding site, type of binding principle (lock-and-key vs. induced fit), ligand solubility, ligand size and speed of diffusion (solute viscosity), all of which are overall not straight forward. A further set of challenges originates from sample delivery: crystals tend to not be perfectly suspended in the microcrystalline slurry, especially at lower solution viscosities and when the crystals are larger than about a micrometre. There have been solutions to this crystal settling problem (Lomb et al., 2012), but those require longer fluidic lines, introducing potential sources of error, since the crystals can settle in capillaries as well as stick to their walls and cause clogging of the lines. Automatic sample exchange for crystalline slurries is also not straight forward, whereas for non-crystalline samples established automated sample dispensing systems (‘auto-sampler’), such as used in HPLC systems and at SAXS beamlines, could be used. It would thus be of great benefit to grow the crystals only when needed, right before the sample is introduced into the X-ray focus – just in time. This would solve many of the challenges: crystal damage, crystal soaking as well as sample delivery, and open completely new possibilities in time-resolved structural biology and structure- or fragment-based drug discovery. To that end, we developed a method for crystallization on-the-fly using the CFEL TapeDrive (Beyerlein et al., 2017; Zielinski et al., 2022) which yields crystals just in time for easy structure determination of biological macromolecules or JINXED (<u>J</u>ust <u>in</u> time <u>c</u>rystallisation for <u>e</u>asy structure <u>d</u>etermination). We present here the first structures obtained with the JINXED method at four different crystallization time points.

## Methods

### Protein sample and crystallizing solution

Hen egg-white lysozyme (Sigma-Aldrich) was prepared at 126 mg/mL in 50 mM acetate buffer pH 3.5. The crystallizing agent contained 0.1 M sodium acetate, pH 4.6, 2.7 M NaCl, 15 % (w/v) PEG4000, 6 % (v/v) ethylene glycol.

### TapeDrive Nozzle

For the first TapeDrive prototype (Beyerlein et al., 2017; Zielinski et al., 2022), sample was deposited onto the tape using polished fused silica fibres with inner diameters ranging from 50 to 180 μm and outer diameters of 360 μm. For mixing on the tape, nozzle-in-nozzle assemblies, derived from developments for double flow-focusing nozzles (Oberthuer et al., 2017), were used. These assemblies were made by fitting a fused silica capillary with an outer diameter smaller than the inner diameter of the second capillary into the second capillary (Wang et al., 2014). A three-way connector (IDEX) was used to decouple outer and inner capillary. Using this connector, the relative position of the end of the inner capillary towards the opening of the outer capillary could be adjusted and thus the contact time between sample and substrate before probing the mixture with X-rays for time-resolved experiments (Beyerlein et al., 2017). The manufacturing of this assembly was not only tedious, but also error prone. Nano-precision 3D-printing of nozzles for liquid jet injection has been established in the past few years and is now being widely used (Knoška et al., 2020; Nelson et al., 2016; Wiedorn et al., 2018). For this study, where mixing of protein solution and crystallizing agent at defined time-steps was required, the design for a 3D-printed gas dynamic virtual nozzles (GDVN, Knoška et al., 2020) was modified and optimized for deposition of sample onto a running tape and simultaneous mixing with a second solution (in this case the crystallizing agent). Since gas-focusing is not needed for sample deposition on the tape, the gas channel was replaced by the mixing channel and mixing on the tape was conducted as in the fast-mixing setting of the original TapeDrive (Beyerlein et al., 2017). The adapted and modified design was subsequently optimized for nano-precision 3D-printing (NanoScribe), resulting in the TapeDrive nozzle (TDN, Figure 2b) as used in this study. However, the usage of the TDN is not limited to the JINXED method but can be used both in mixing and non-mixing mode for sample delivery with the CFEL TapeDrive. Depending on tape speed and TDN position relative to the X-ray focus, time delays between 50 ms and 100 s can be realised with this set-up. The TDN has a size of 970 μm x 450 μm x 860 μm (l x w x h, maximum dimensions from bottom to tip) and the orifice of the combined sample and mixing channel is 200 μm wide. The two fused silica capillaries feeding sample and crystallizing solution into the nozzle have an inner diameter of 150 μm.

**Figure 1:**
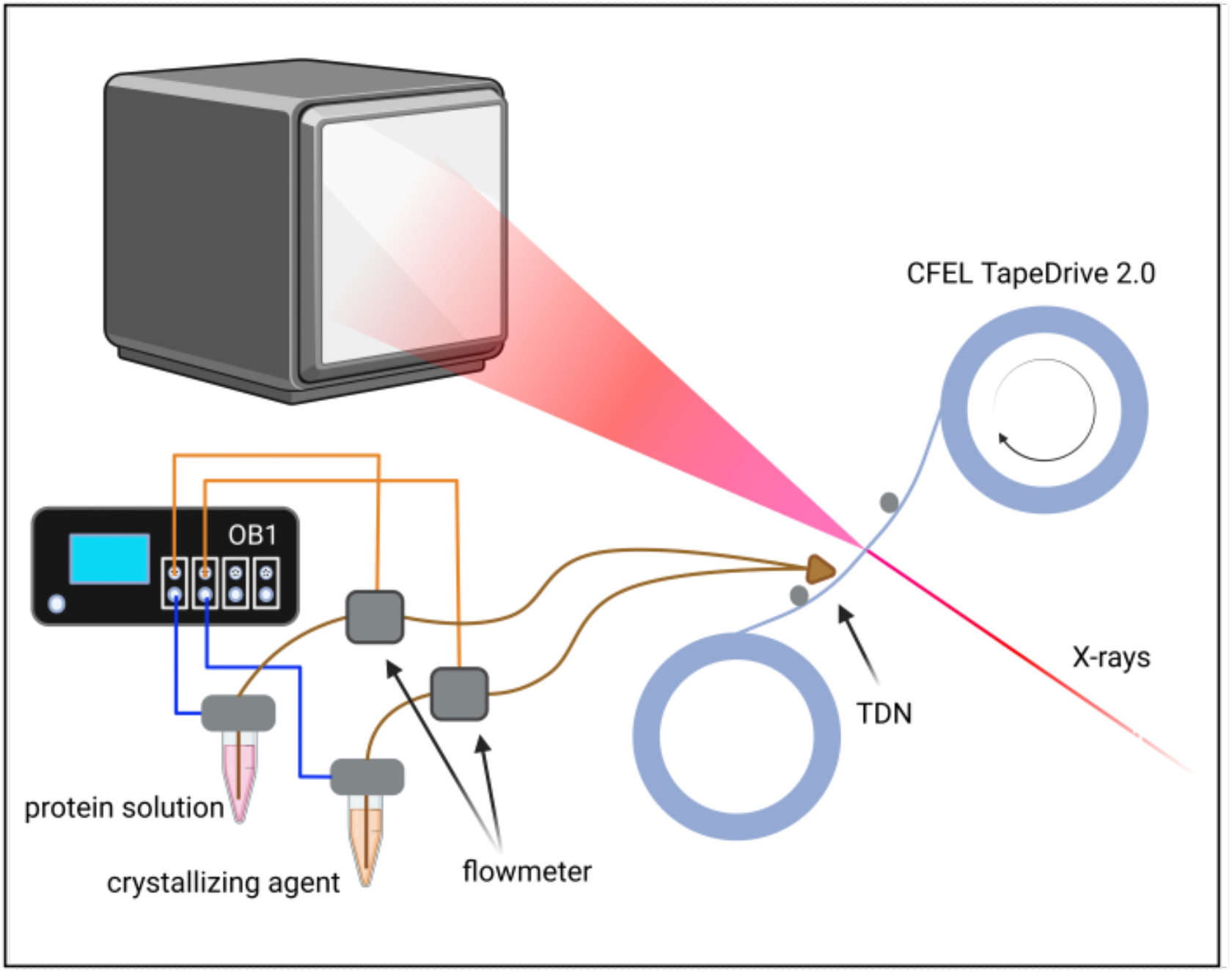
schematic drawing of experimental setup showing the Elveflow controller OB1, microfluidic flow rate sensors (flowmeters), reservoirs with protein solution and crystallization buffer, the sample delivery system CFEL TapeDrive 2.0 including the TapeDrive nozzle (TDN), X-ray beam and detector Eiger2 × 16M.

**Figure 2:**
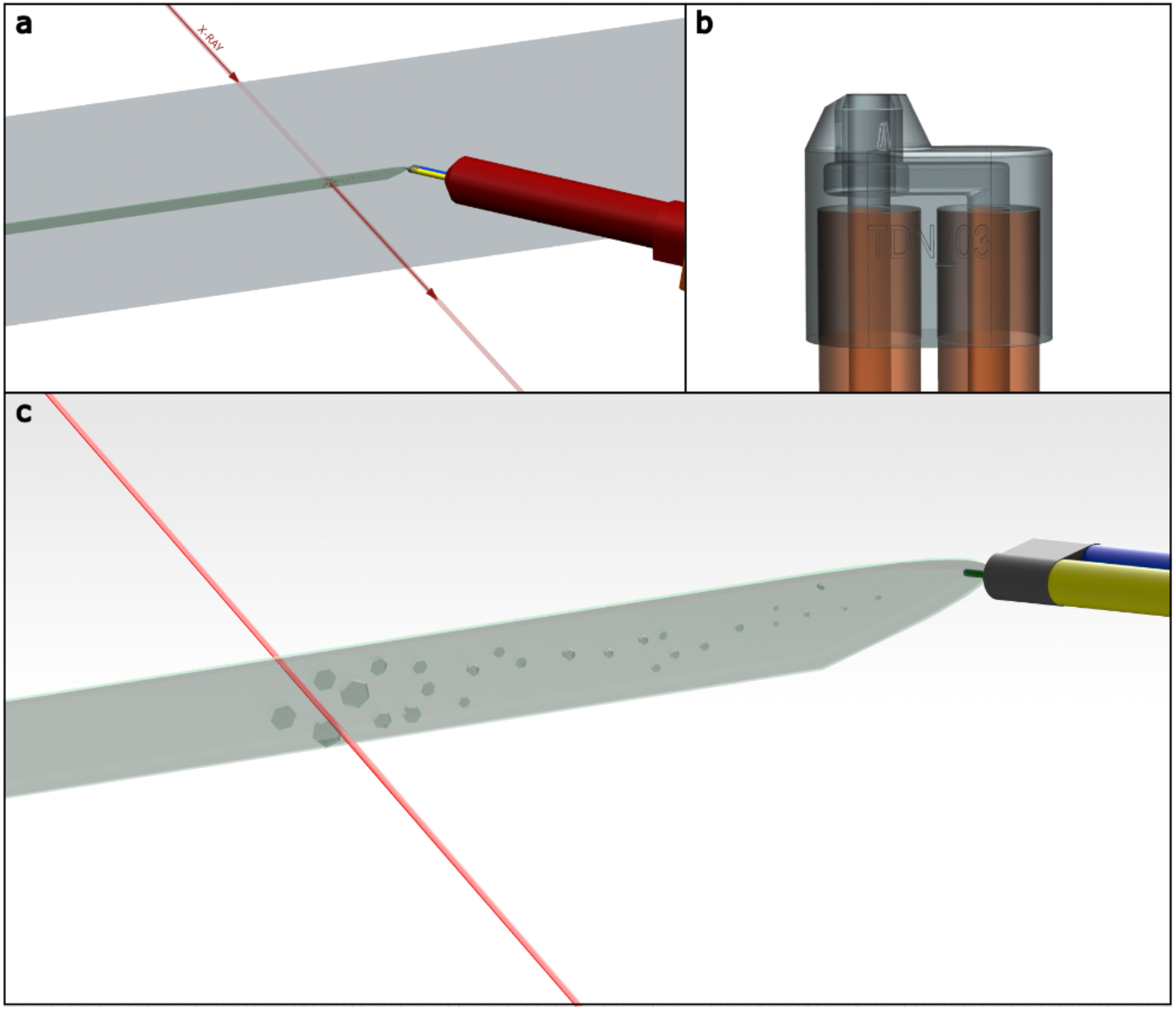
Drawings of a) sample environment overview with TapeDrive nozzle, tape, sample line and X-ray beam, b) TapeDrive nozzle with inner (left) and outer mixing (right) channel, c) JINXED principle with TapeDrive nozzle incorporating the protein solution channel (yellow) and crystallizing agent channel (blue). Due to mixing within the sample line on the tape, protein crystallization can be observed.

### Data collection at P11 using the JINXED method

Sample delivery was performed by the CFEL TapeDrive 2.0 (manuscript in preparation), an updated version of the CFEL TapeDrive (Beyerlein et al., 2017). The general description of sample delivery using a microfluidic controller as described (Zielinski et al., 2022) is still valid for TapeDrive 2.0 (see Figure 1 and Figure 2a) that was optimized for fast installation at beamlines, ease of use, low sample consumption as well as more accurate tape movement. For the JINXED method this was just slightly modified: the protein solution was placed in a reservoir and transported to the TDN using an Elveflow OB1 flow controller. The reservoir was connected directly to this controller and pressurized by the controller and not indirectly through a plunger. A Microfluidic Flow Rate Sensor (Elveflow, France) was placed in line after the reservoir to give feedback on the flow rate corresponding to a certain input pressure. The nozzle, as described above, was connected to the microfluidic tubing using standard HPLC-connectors (IDEX, USA). For the JINXED experiment, a second reservoir, containing the crystallizing agent, was connected to the nozzle in the same way, using a different channel at the OB1 flow controller for independent control of both flow rates. The reservoirs, standard micro reaction tubes with either 1.5 mL or 15 mL volumes can be placed in a heating/cooling block or in a shaker for such tubes (e.g. Eppendorf ThermoMix C) for prolonged sample integrity. Usage of fused silica fibres (Polymicro, USA) with an inner diameter of 150 μm as a connection between the 3D-printed nozzle and the sample reservoirs, ensured a smooth sample flow with as little restriction and clogging probability as possible. Protein and crystallizing solution were mixed in a 1:1 ratio at a flow rate of 1 μL/min each, resulting in protein crystallization (Figure 2c, supplementary Figure 6). The nozzle-beam distance was varied from 2 - 8 mm (at a tape speed of 1 mm s^−1^) to probe crystallization times of 2 s, 4 s, 6 s and 8 s. X-ray data collection was carried out at beamline P11 (PETRA III, DESY, Hamburg) using 12.0 keV photon energy X-rays focused to a spot of 4 × 9 μm (width × height) with a flux of 8.6 × 10^12^ photons s^−1^. *Raddose-3D* (Zeldin et al., 2013) was used to estimate the dose, assuming a crystal size range of 1 x 1 x 1 μm to 5 x 5 x 5 μm size. The script given in the supplementary material resulted in a dose range of 0.23 MGy to 0.30 MGy.

Data were continuously collected for 1 h for each data set, using an EIGER2 × 16M detector at 130 Hz frame rate, providing an exposure time of 7.69 ms. Feedback about hit und indexing rate of the incoming data was given by the *OnDA* software package (Mariani et al., 2016).

### Data processing

Raw data files were processed with *CrystFEL 0.9.1* (White et al., 2012, 2016) using custom scripts. In *indexamajig*, the option --peaks=peakfinder8 was used to identify individual ‘hits’ from the complete set of collected diffraction patterns. The complete set was defined by an automatically generated list of files. Detected ‘hits’ were then indexed using XGANDALF (Gevorkov et al., 2019) and integrated. The geometry input file was adapted for the photon energy and detector distance from previous experiments at P11. The resulting stream-files were merged into point group 4/mmm using *partialator* and figures of merit calculated using *compare_hkl* and *check_hkl,* all part of the CrystFEL package. MTZ files for crystallographic data processing were generated from CrystFEL merged reflection datafiles using F2MTZ within the CCP4 suite (Winn et al., 2011).

Starting model for refinement was the lysozyme structure with PDB accession code 6FTR (Wiedorn et al., 2018) for all data sets and Phenix (Liebschner et al., 2019) was used for refinement and model validation. *R*free flags were generated using *phenix.reflection_file_editor* and the same set of Rfree flags was used for all datasets in this study. In *phenix.refine* (Afonine et al., 2012) B-factors of the starting model were set to 20.0 for the first round of refinement, followed by visual inspection of the model and maps using *Coot* (Emsley et al., 2010). Further iterative cycles of refinement included TLS, optimization of X-ray/stereochemistry and X-ray/ADP weights as well as manual model building in *Coot*. *MolProbity* (Williams et al., 2017) was used for validation of the final model. Figures were generated using PyMOL, which was also used for alignment of structures and r.m.s.d. calculations.

## Results

Diffraction from protein crystals was observed first at 2 s after mixing of protein solution and crystallizing agent at a nozzle to X-ray focus distance of 2 mm and a tape speed of 1 mms^−1^, manifesting the proof-of-principle of simultaneous protein crystallization and diffraction data collection, named JINXED – <u>J</u>ust <u>in</u> time <u>c</u>rystallization for <u>e</u>asy structure <u>d</u>etermination. At a mixing time of 2 s, 40.6 % of all recorded detector frames contained a ‘hit’ and 21.1 % of those hits contained indexable patterns. After 4 s of mixing, at a nozzle to X-ray focus distance of 4 mm and at a tape speed of 1 mm s^−1^ the hit rate increased to 43.6 %, whereas the indexing rate decreased to 18.6 %. A mixing time of 6 s (corresponding to a distance of 6 mm) resulted in a hit rate of 83.1 % and an indexing rate of 42.1 %. Extending the crystallization time to 8 s (8 mm nozzle to X-ray focus distance) led to an increase of the hit rate to 94.8 % and a decrease of the indexing rate to 25.9 %. Detailed data collection and refinement statistics can be found in table 1.

**Table 1.**
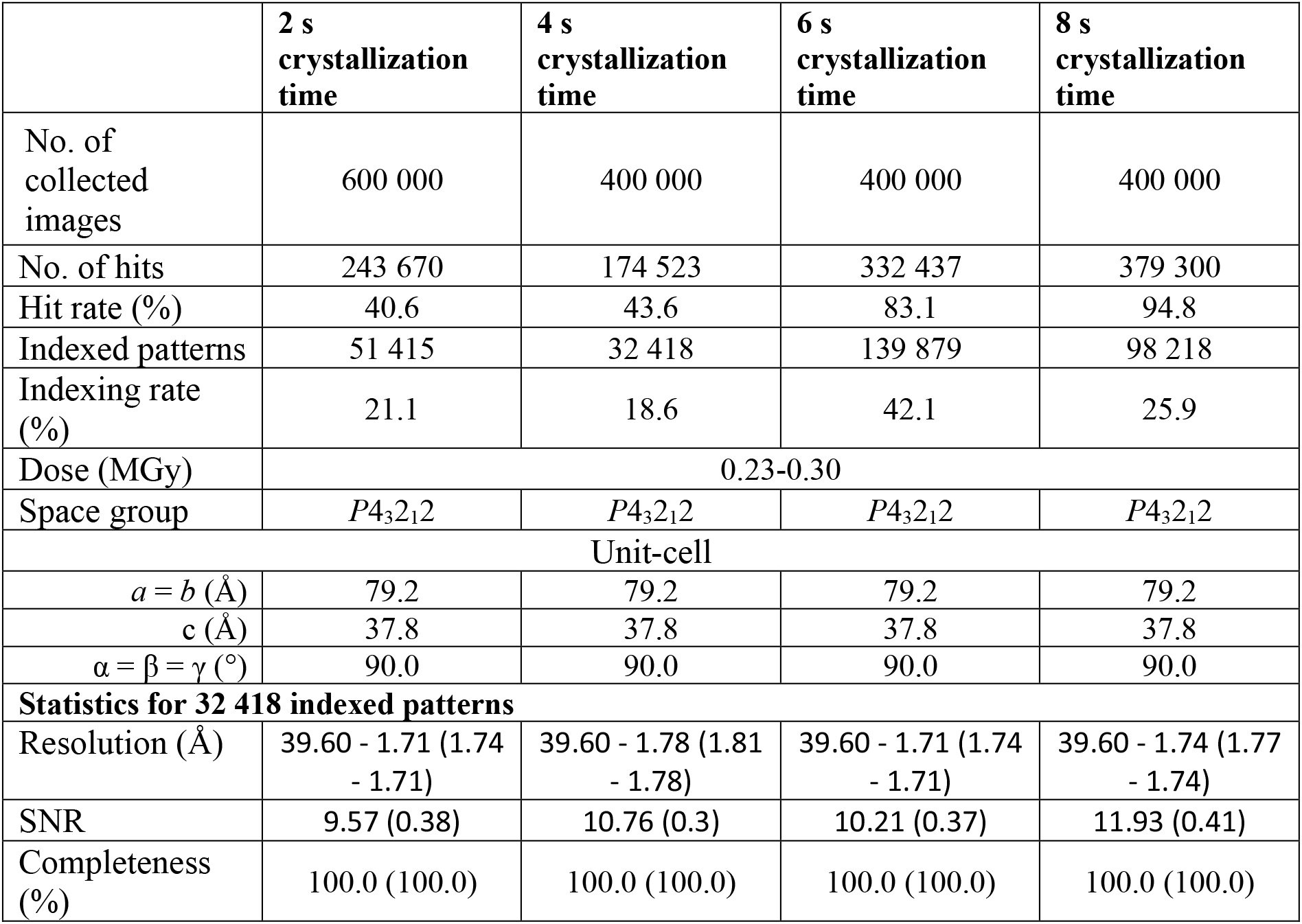
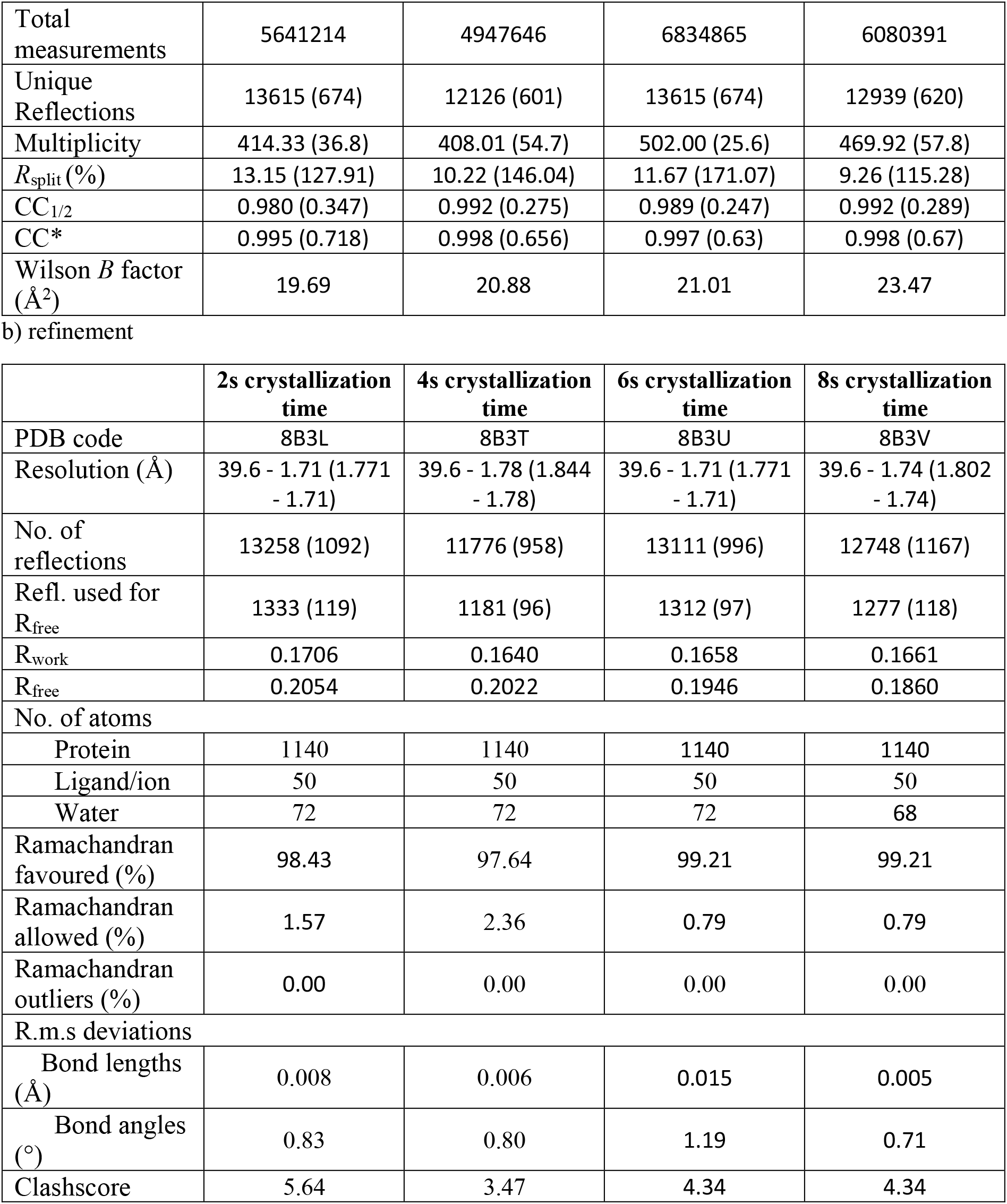

For the dataset recorded at 4 s crystallization time, a total of 32 418 indexed lattices could be found. For better comparability of the four data sets, all were cut to 32 418 randomly selected indexed lattices prior to calculating figures of merit (Table 1). The resolution cut-off was chosen individually for each data set corresponding to a CC* value > 0.5. Between the four data sets, the resolution differs only by 0.07 Å. The overall SNR increases slightly with increasing crystallization time from 9.57 (2 s) to 10.76 (4 s), 10.21 (6 s) and 11.93 (8 s).

As shown in Figure 3, all data sets yielded 2Fo-Fc electron density maps of very similar quality and no clear trend between map quality and crystallization time could be observed. Refinement (from starting model with the PDB accession code 6FTR, Wiedorn et al., 2018) resulted in R_work_ values of 0.171 (2 s), 0.164 (4 s), 0.166 (6 s), and 0.166 (8 s) with no clear tendency along the increasing crystallization time. R_free_ values decreased slightly with increasing crystallization time from 0.205 (2 s), 0.202 (4 s), 0.195 (6 s) to 0.186 (8 s). The refined structures of all four crystallization times were aligned with negligible RMSD (< 0.05). Depicted in Figure 4 is the overlay of all structure models (4a), the overlay of the two residues Glu35 and Asp52, forming the active site (4b), and the active site overlay of the 8 s model and that from 6FTR (4c), showing no major differences between the structural models.

**Figure 3:**
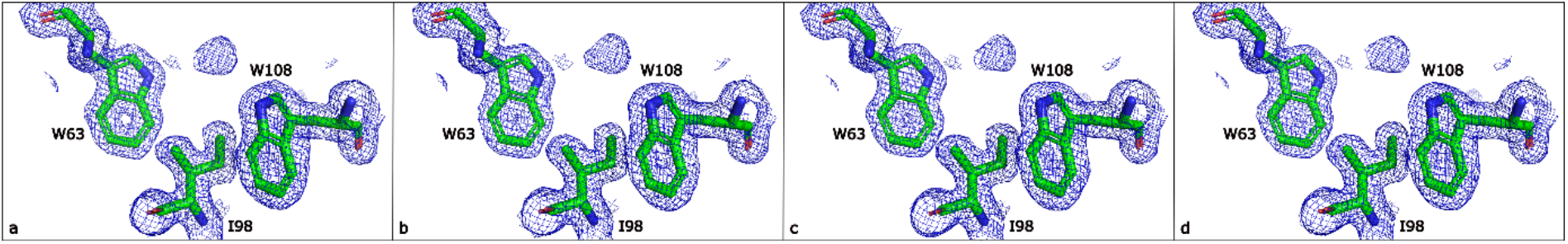
electron-density maps (contour level 1.0) with models of residues W63, I98, W108 of a) 2 s crystallization time, b) 4 s crystallization time, c) 6 s crystallization time, and d) 8 s crystallization time

**Figure 4:**
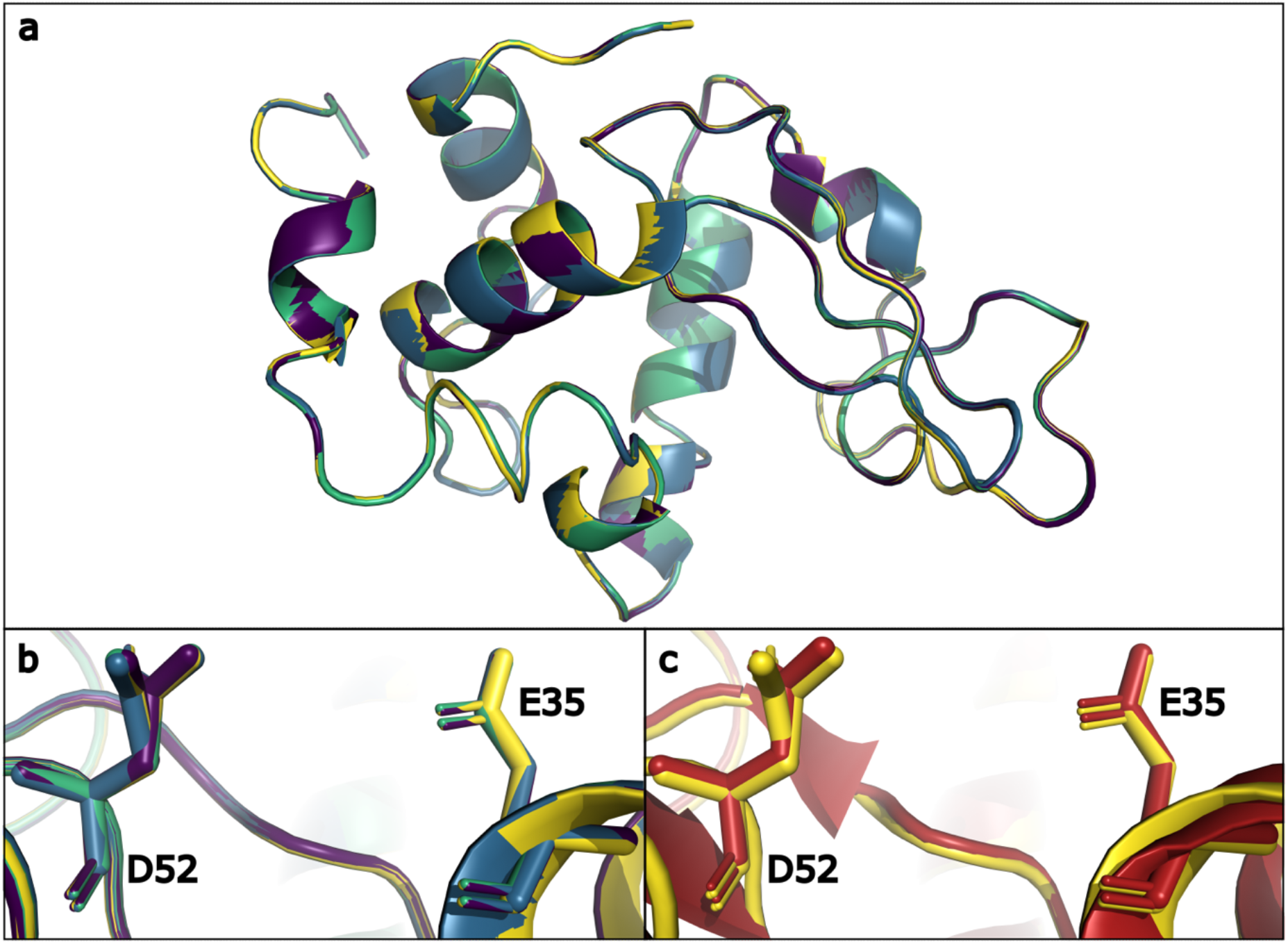
Structure overlay of a) structures from all data sets with 2 s (violet), 4 s (blue), 6 s (green), and 8 s (yellow) crystallization time; b) residues D52 and E35 from all structures from all data sets with 2 s (violet), 4 s (blue), 6 s (green), and 8 s (yellow) crystallization time; c) residues D52 and E35 from data set with 8 s crystallization (yellow) and 6ftr model (red).

## Conclusion

The experiments presented here using the JINXED method are the first of their kind and therefore provide a proof-of-principle for online crystallization and diffraction. The underlying idea was developed based on the crystallization behaviour of hen egg white lysozyme (HEWL), which forms crystals immediately after mixing with the crystallizing agent, and the possibility for rapid mix-and-diffuse experiments with the TapeDrive setup (Beyerlein et al., 2017). Diffraction from the HEWL microcrystals was detected just two seconds after mixing protein and crystallizing agent, showing that HEWL forms crystals of sufficient size for SSX at a 3^rd^ generation synchrotron within this time frame.

The JINXED method has the advantage that the sensitive microcrystals do not need to be handled in any form prior to data collection. This eliminates the occurrence of crystal damage caused by mechanical stress, which is reported to often limit the diffraction resolution of crystals (Dobrianov *et al.*, 1999).

Regarding the variety of crystallographic experiments, even time-resolved mix-and-diffuse experiments or inhibitor and fragment binding studies can be conducted within our approach. This has the potential to solve the problem of insufficient diffusion saturation of the crystals due to various reasons (see introduction), which is faced by other data collection methods of serial time-resolved studies. Furthermore, it is less tedious than co-crystallization prior to the experiment. In principle, compounds for structure- or fragment-based drug design could be incubated with protein in e.g. 384 well SBS-format plates and an auto-sampler such as used in HPLC instruments could then dispense the protein-compound solutions one after the other to the TDN for JINXED and high-throughput data collection using the TapeDrive 2.0 (see Figure 5). Similarly, for time-resolved enzyme studies, substrate or analogues could be mixed with crystallizing agent. The resulting time delay would be equal to that of the crystallization time. However, at XFELs crystals with much smaller size (few 100 nm in diameter, Gati et al., 2017) can be used and thus crystals of sufficient quality could be available in a few ms. In addition to mix-and-diffuse, it would also be compatible with optical reaction triggering and pH-jump experiments.

**Figure 5:**
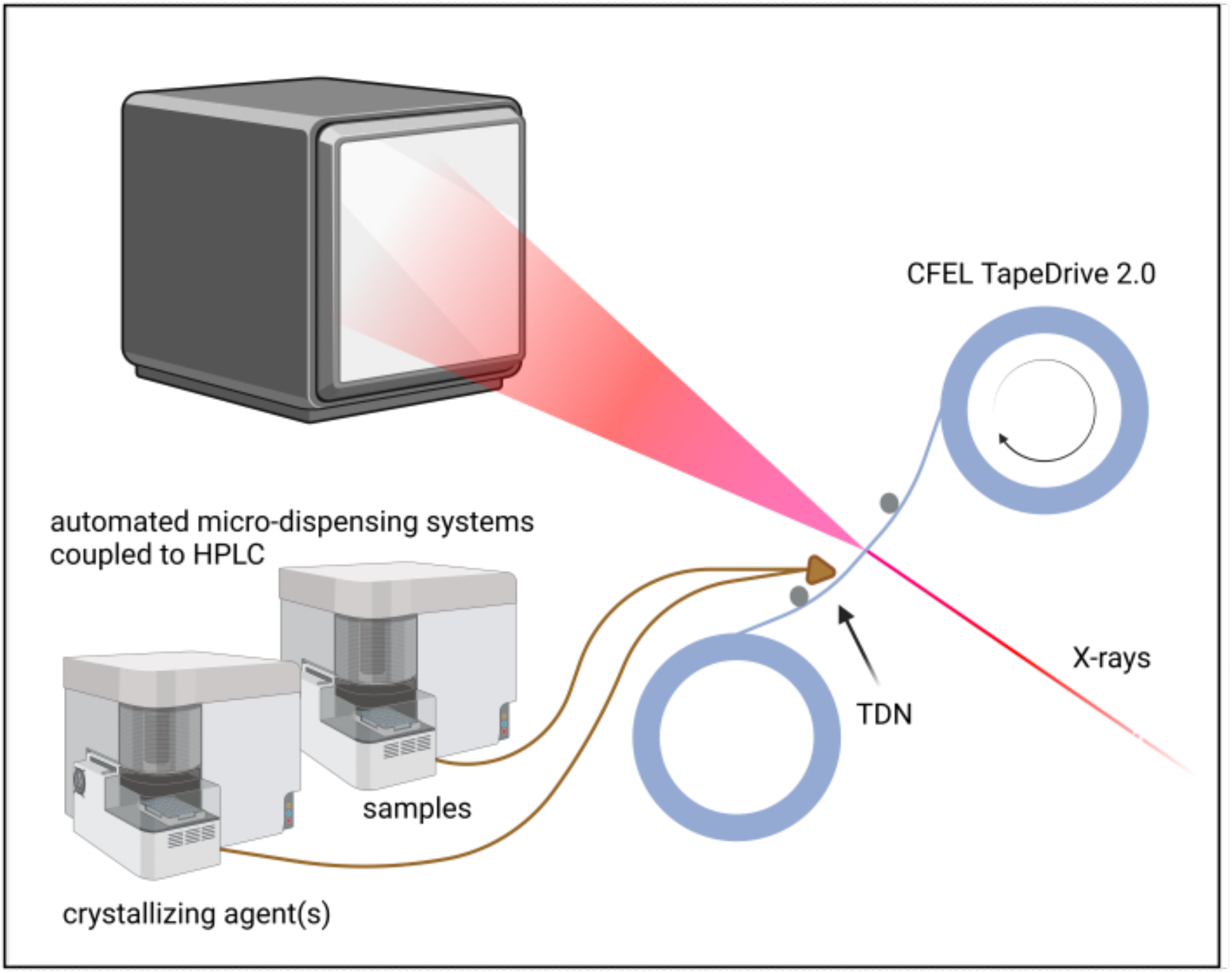
schematic drawing of a possible high-throughput setup showing two automated micro-dispensing systems for samples (e.g. protein mixed with compound) and crystallization agents, the sample delivery system CFEL TapeDrive 2.0 including the TapeDrive nozzle (TDN), X-ray beam and detector.

Nonetheless, JINXED experiments require fast crystallization of the sample. The screening for suitable crystallization conditions may be time-consuming and extensive, but this challenge is equally faced in protein crystallography in general (Chayen, 2004; Chayen and Saridakis, 2008).Once successful crystallization conditions are determined, optimization of the latter according to the general phase behaviour of proteins could lead to faster crystallization rates suitable for this approach. The focus of protein crystallization research has been, obviously, on the reliable production of large, well diffracting crystals, with few examples focussing on the directed generation of small crystals for solid state NMR (Martin and Zilm, 2003), SFX and SSX (Beale et al., 2019; Stohrer et al., 2021; Tenboer et al., 2014). The JINXED method could even be used to screen for crystallization conditions and crystallization times without the need for visual assessment and direct proof of crystalline properties. It can be assumed that many proteins could be crystallized rapidly if conditions would be optimized towards velocity of crystallization as reported in Santarsiero et al., 2002. However, a further understanding of the underlying physics of protein nucleation and crystallization and further research in that direction is necessary, but way beyond the scope of this study. Again, the adaption to X-ray free-electron laser beamlines would lower the required crystal size, which may enable JINXED experiments for proteins that show slower growth rates. For SFX experiments it would be beneficial to use a sample delivery method that avoids the introduction of superfluous material to the beam, since this occurs to be problematic for the high-intensity X-ray pulses produced by XFELs. For example, liquid jets generated by nozzles with integrated mixing devices (Knoška et al., 2020) may be suitable for implementing the JINXED method at XFELs.

The variety of proteins suitable for JINXED experiments at SSX beamlines can be broadened by exploiting the TapeDrive’s possibility for prolonged mixing/crystallization times of up to several minutes (Beyerlein et al., 2017), which improves the general applicability of this method. Moreover, the addition of temperature control to the TapeDrive might facilitate rapid crystallization for proteins whose solubility is temperature dependent. On that note, recently multi-dimensional studies became apparent in serial crystallography to investigate structural changes as response to temperature changes (Mehrabi et al., 2021). Here, the experiment could again benefit from the strategy presented in this study, since the crystals are directly grown in the different environments and not confronted with environment changes after growth, reducing stress on the crystal lattice and thus potentially increasing diffraction quality.

Due to the numerous advantages of the JINXED approach as outlined above, this method should be further evaluated, including different proteins, setups, and beamlines to assess the overall applicability of this novel approach. Even if the method is found to be suitable for a subset of proteins of biological or pharmaceutical relevance, it could be a game changer for high-output studies of enzyme dynamics and drug- and fragment-binding properties.

## Acknowledgements

This research was supported in part through the Maxwell computational resources operated at Deutsches Elektronen-Synchrotron DESY, Hamburg, Germany. We acknowledge DESY (Hamburg, Germany), a member of the Helmholtz Association HGF, for the provision of experimental facilities. Parts of this research were carried out at PETRA III, beamline P11, and we would like to thank Eva Crosas, Jan Meyer, Helena Taberman and Guillaume Pompidor for assistance with the experiment. This work was funded by Deutsche Forschungsgemeinschaft (DFG, German Research Foundation) – 491245950. Some images were created with https://Biorender.com.

## Competing interests

The authors declare no competing interests.

## Supplementary

**FigureS6:**
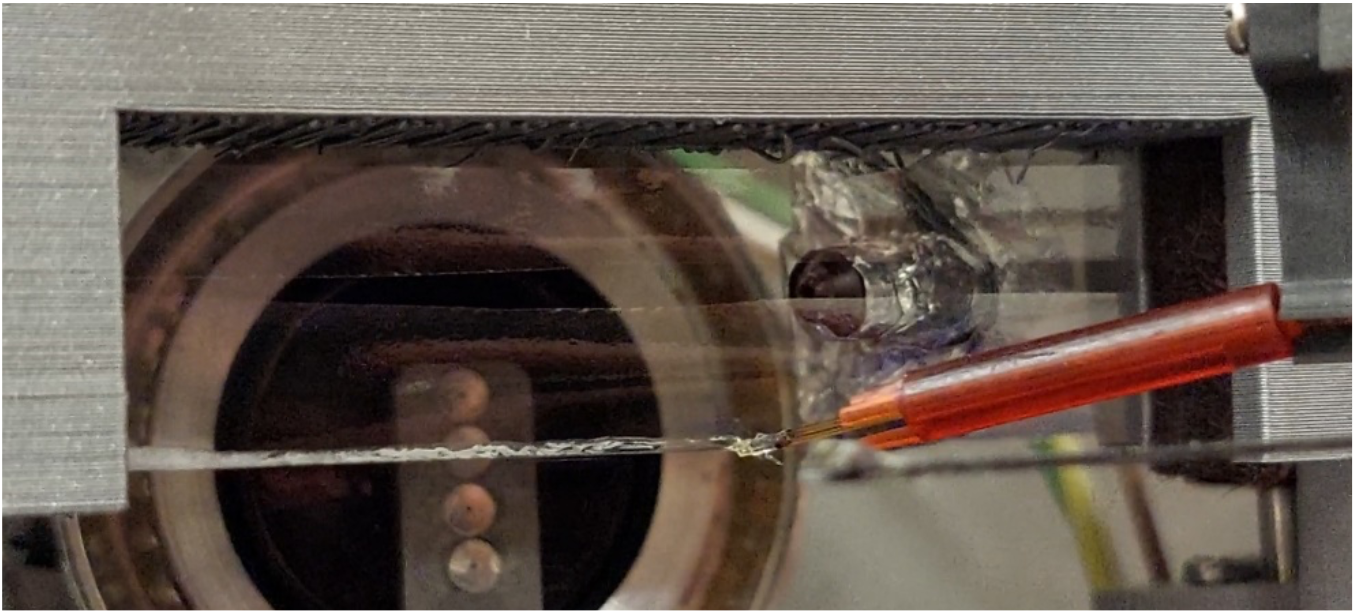
photo of JINXED at P11, DESY, HH. The 3D-printed TapeDrive nozzle deposits the protein solution and crystallizing agent onto the tape where mixing occurs subsequently. The white clouding within the sample line indicates protein crystallization.

---

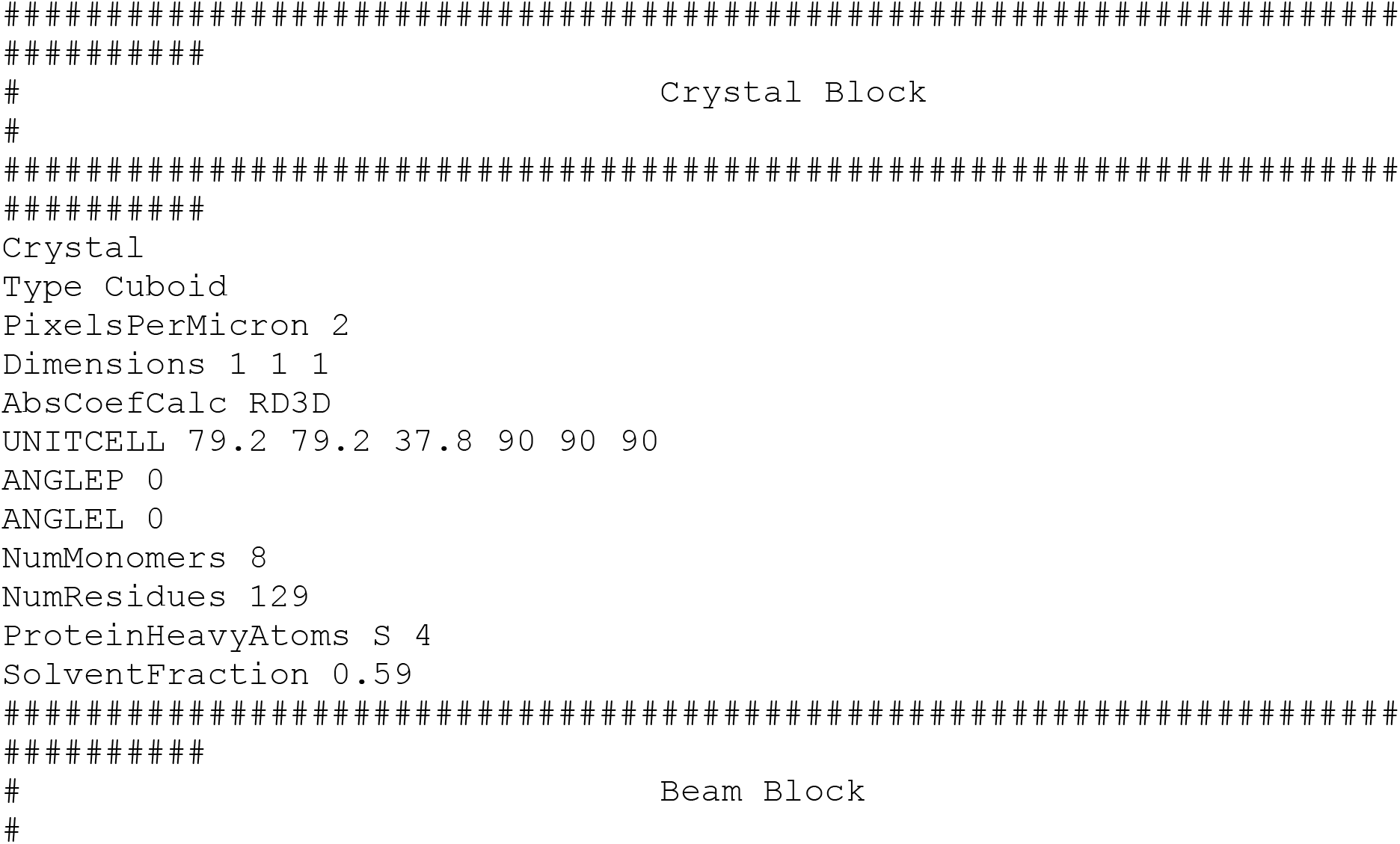
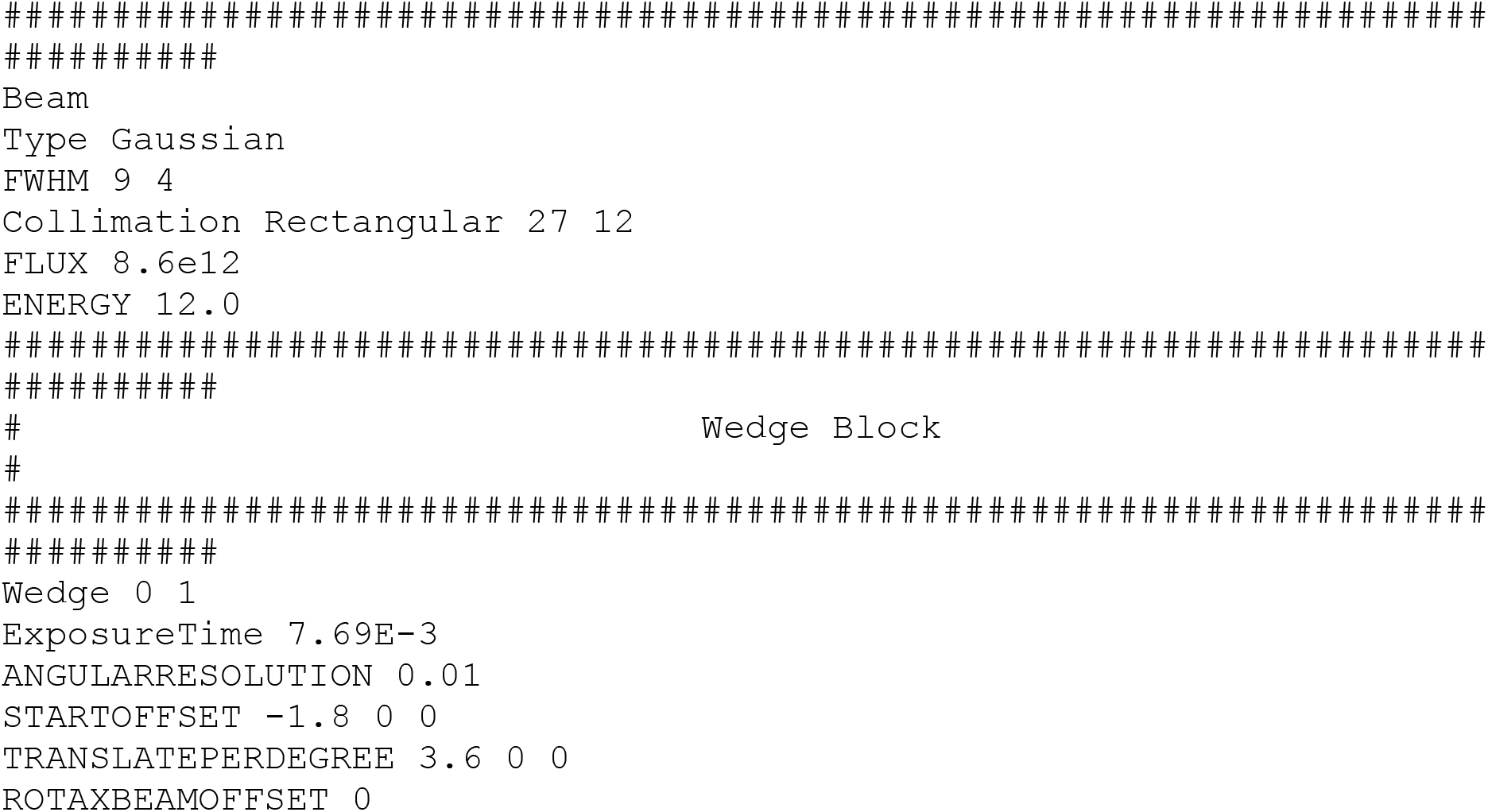
Raddose-3D script for the X-ray dose calculation for 1×1×1 μm crystals.

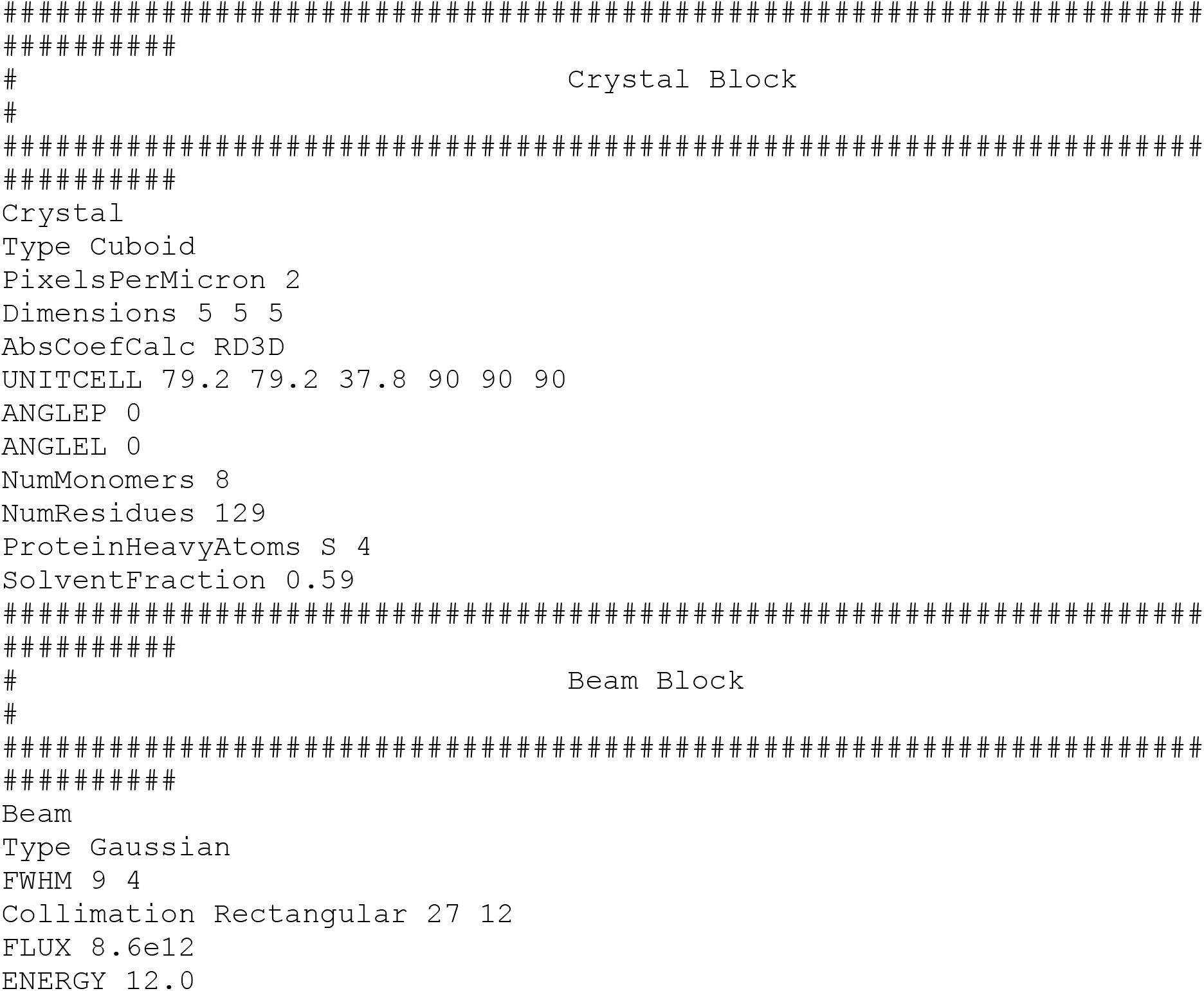
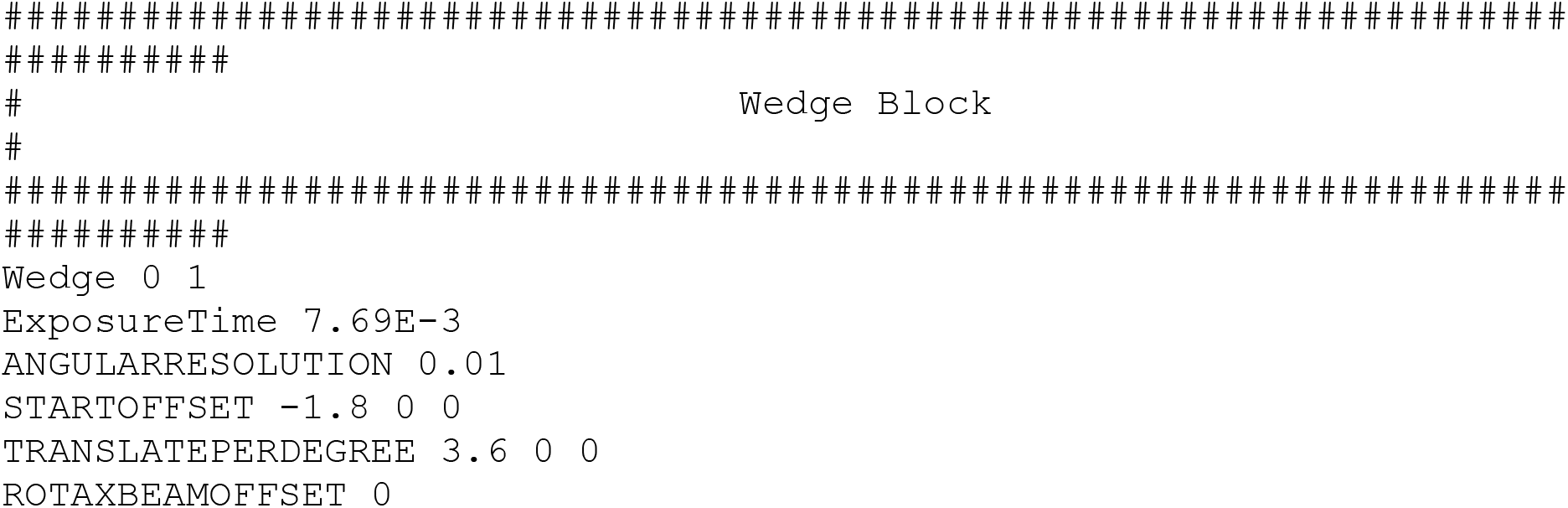
Raddose-3D script for the X-ray dose calculation for 5×5×5 μm crystals.

## Notes

### Competing Interest Statement

The authors have declared no competing interest.

### Summary of Updates

A citation error was corrected and an accepted manuscript that was cited got published and citation was updated.

## References

Afonine, P. V., Grosse-Kunstleve, R.W., Echols, N., Headd, J.J., Moriarty, N.W., Mustyakimov, M., Terwilliger, T.C., Urzhumtsev, A.G., Zwart, P.H., and Adams, P.D. (2012). Towards automated crystallographic structure refinement with phenix. refine research papers. Acta Crystallogr. - Sect. D Biol. Crystallogr. 68, 352–367.

Barends, T.R.M., Stauch, B., Cherezov, V., and Schlichting, I. (2022). Serial femtosecond crystallography. Nat Rev Methods Prim. 2 (59).

Beale, J.H., Bolton, R., Marshall, S.A., Beale, E. V., Carr, S.B., Ebrahim, A., Moreno-chicano, T., Hough, M.A., Worrall, J.A.R., Tews, I., et al. (2019). Successful sample preparation for serial crystallography experiments. J. Appl. Crystallogr. 52 (6), 1385–1396.

Beyerlein, K.R., Dierksmeyer, D., Mariani, V., Kuhn, M., Sarrou, I., Ottaviano, A., Awel, S., Knoska, J., Fuglerud, S., Jönsson, O., et al. (2017). Mix-and-diffuse serial synchrotron crystallography. IUCrJ 4, 769–777.

Boutet, S., Lomb, L., Williams, G.J., Barends, T.R.M., Aquila, A., Doak, R.B., Weierstall, U., DePonte, D.P., Steinbrener, J., Shoeman, R.L., et al. (2012). High-Resolution Protein Structure Determination by Serial Femtosecond Crystallography. Science (. 337, 362–364.

Brändén, G., and Neutze, R. (2021). Advances and challenges in time-resolved macromolecular crystallography. Science. 373 (6558).

Chapman, H.N., Fromme, P., Barty, A., White, T.A., Kirian, R.A., Aquila, A., Hunter, M.S., Schulz, J., Deponte, D.P., Weierstall, U., et al. (2011). Femtosecond X-ray protein nanocrystallography. Nature 470, 73–78.

Chayen, N.E. (2004). Turning protein crystallisation from an art into a science. Curr Opin Struct Biol. 5, 577–583.

Chayen, N.E., and Saridakis, E. (2008). Protein crystallization: from purified protein to diffraction-quality crystal. 5, 147–153.

Dickerson, E. (2005). Present at the Flood: How Structural Molecular Biology Came About, Richard E. Dickerson (2005), Sinauer Associates, Sunderland, Massachusetts. FASEB J. 20, 809–810.

Dobrianov, I., Caylor, C., Lemay, S.G., Finkelstein, K.D., and Thorne, R.E. (1999). X-ray diffraction studies of protein crystal disorder. J. Cryst. Growth 196, 511–523.

Emsley, P., Lohkamp, B., Scott, W.G., and Cowtan, K. (2010). Features and development of Coot. Acta Crystallogr. Sect. D Biol. Crystallogr. 66, 486–501.

Gati, C., Oberthuer, D., Yefanov, O., Bunker, R.D., Stellato, F., and Chiu, E. (2017). Atomic structure of granulin determined from native nanocrystalline granulovirus using an X-ray free-electron laser. PNAS 114, 2247–2252.

Gevorkov, Y., Yefanov, O., Barty, A., White, T.A., Mariani, V., Brehm, W., Tolstikova, A., Grigat, R.R., and Chapman, H.N. (2019). XGANDALF - Extended gradient descent algorithm for lattice finding. Acta Crystallogr. Sect. A Found. Adv. 75, 694–704.

Giegé, R. (2013). A historical perspective on protein crystallization from 1840 to the present day. FEBS J. 280, 6456–6497.

Hansen, C., and Quake, S.R. (2003). Microfluidics in structural biology: Smaller, faster... better. Curr. Opin. Struct. Biol. 13, 538–544.

Heymann, M., Opthalage, A., Wierman, J.L., Akella, S., Szebenyi, D.M.E., Gruner, S.M., and Fraden, S. (2014). Room-temperature serial crystallography using a kinetically optimized microfluidic device for protein crystallization and on-chip X-ray diffraction. IUCrJ 1, 349–360.

Knoška, J., Adriano, L., Awel, S., Beyerlein, K.R., Yefanov, O., Oberthuer, D., Peña Murillo, G.E., Roth, N., Sarrou, I., Villanueva-Perez, P., et al. (2020). Ultracompact 3D microfluidics for time-resolved structural biology. Nat. Commun. 11, 1–12.

Liebschner, D., Afonine, P. V., Baker, M.L., Bunkóczi, G., Chen, V.B., Croll, T.I., Hintze, B., Hung, L.-W., Jain, S., McCoy, A.J., et al. (2019). Macromolecular structure determination using X-rays, neutrons and electrons: recent developments in Phenix. Acta Crystallogr. - Sect. D Biol. Crystallogr. 861–877.

Lomb, L., Steinbrener, J., Bari, S., Beisel, D., Berndt, D., Kieser, C., Lukat, M., Neef, N., and Shoeman, R.L. (2012). An anti-settling sample delivery instrument for serial femtosecond crystallography research papers. J. Appl. Crystallogr. 45, 674–678.

Mariani, V., Morgan, A., Yoon, H., Lane, T.J., White, T.A., Grady, C.O., Kuhn, M., and Aplin, S. (2016). OnDA: online data analysis and feedback for serial. J. Appl. Crystallogr. 49 (3), 1073–1080.

Martin-Garcia, J.M. (2021). Protein dynamics and time resolved protein crystallography at synchrotron radiation sources: Past, present and future. Crystals 11.

Martin, R.W., and Zilm, K.W. (2003). Preparation of protein nanocrystals and their characterization by solid state NMR. J. Magn. Reson. 165, 162–174.

McPherson, A. (1991). A brief history of protein crystal growth. J. Cryst. Growth 110, 1–10.

McPherson, A., and Gavira, J.A. (2014). Introduction to protein crystallization. Acta Crystallogr. Sect. FStructural Biol. Commun. 70, 2–20.

Mehrabi, P., Schulz, E.C., Dsouza, R., Müller-Werkmeister, H.M., Tellkamp, F., Dwayne Miller, R.J., and Pai, E.F. (2019). Time-resolved crystallography reveals allosteric communication aligned with molecular breathing. Science. 365, 1167–1170.

Mehrabi, P., von Stetten, D., Leimkohl, J.-P., Tellkamp, F., and Schulz, E.C. (2021). An environmental control box for serial crystallography enables multi-dimensional experiments. BioRxiv 2021.11.07.467596.

Nelson, G., Kirian, R.A., Weierstall, U., Zatsepin, N.A., Baumbach, T., Wilde, F., Niesler, F.B.P., Zimmer, B., Ishigami, I., Hikita, M., et al. (2016). Three-dimensional-printed gas dynamic virtual nozzles for x-ray laser sample delivery. Opt. Express 24, 11515–11530.

Oberthuer, D., Knoška, J., Wiedorn, M.O., Beyerlein, K.R., Bushnell, D.A., Kovaleva, E.G., Heymann, M., Gumprecht, L., Kirian, R.A., Barty, A., et al. (2017). Double-flow focused liquid injector for efficient serial femtosecond crystallography. Sci. Rep. 7.

Pande, K., Hutchison, C.D.M., Groenhof, G., Aquila, A., Robinson, S., Tenboer, J., Basu, S., Boutet, S., Deponte, D.P., Liang, M., et al. (2016). Femtosecond Structural Dynamics Drives the Trans/Cis Isomerization in photoactive Yellow Protein. HHS Public Access 352, 725–729.

Perry, S.L., Guha, S., Pawate, A.S., Bhaskarla, A., Agarwal, V., Nair, S.K., and Kenis, P.J.A. (2013). A microfluidic approach for protein structure determination at room temperature via on-chip anomalous diffraction. Lab Chip 13, 3183–3187.

Santarsiero, B.D., Yegian, D.T., Lee, C.C., Spraggon, G., Gu, J., Scheibe, D., Uber, D.C., Cornell, E.W., Nordmeyer, R.A., Kolbe, W.F., et al. (2002). An approach to rapid protein crystallization using nanodroplets. J. Appl. Crystallogr. 35, 278–281.

Stagno, J.R., Liu, Y., Bhandari, Y.R., Conrad, C.E., Panja, S., Swain, M., Fan, L., Nelson, G., Li, C., Wendel, D.R., et al. (2017). Structures of riboswitch RNA reaction states by mix-and-inject XFEL serial crystallography. Nature 541, 242–246.

Stohrer, C., Horrell, S., Meier, S., Sans, M., von Stetten, D., Hough, M., Goldman, A., Monteiro, D.C.F., and Pearson, A.R. (2021). Homogeneous batch micro-crystallization of proteins from ammonium sulfate. Acta Crystallogr. Sect. D Biol. Crystallogr. 77, 194–204.

Tenboer, J., Basu, S., Zatsepin, N., Pande, K., Milathianaki, D., Frank, M., Hunter, M., Boutet, S., Williams, G.J., Koglin, J.E., et al. (2014). Time-resolved serial crystallography captures high-resolution intermediates of photoactive yellow protein. Science. 346, 1242–1246.

Wang, D., Weierstall, U., Pollack, L., and Spence, J. (2014). Double-focusing mixing jet for XFEL study of chemical kinetics. J. Synchrotron Radiat. 21, 1364–1366.

White, T.A., Kirian, R.A., Martin, A. V., Aquila, A., Nass, K., Barty, A., and Chapman, H.N. (2012). CrystFEL: A software suite for snapshot serial crystallography. J. Appl. Crystallogr. 45, 335–341.

White, T.A., Mariani, V., Brehm, W., Yefanov, O., Barty, A., Beyerlein, K.R., Chervinskii, F., Galli, L., Gati, C., Nakane, T., et al. (2016). Recent developments in CrystFEL. J. Appl. Crystallogr. 49, 680–689.

Wiedorn, M.O., Oberthür, D., Bean, R., Schubert, R., Werner, N., Abbey, B., Aepfelbacher, M., Adriano, L., Allahgholi, A., Al-Qudami, N., et al. (2018). Megahertz serial crystallography. Nat. Commun. 9, 1–11.

De Wijn, R., Hennig, O., Roche, J., Engilberge, S., Rollet, K., Fernandez-Millan, P., Brillet, K., Betat, H., Mörl, M., Roussel, A., et al. (2019). A simple and versatile microfluidic device for efficient biomacromolecule crystallization and structural analysis by serial crystallography. IUCrJ 6, 454–464.

Williams, C.J., Headd, J.J., Moriarty, N.W., Prisant, M.G., Videau, L.L., Deis, L.N., Verma, V., Keedy, D.A., Hintze, B.J., Chen, V.B., et al. (2017). MolProbity: More and better reference data for improved all-atom structure validation. Protein Sci. 27(1), 293–315.

Winn, M.D., Ballard, C.C., Cowtan, K.D., Dodson, E.J., Emsley, P., Evans, P.R., Keegan, R.M., Krissinel, E.B., Leslie, A.G.W., McCoy, A., et al. (2011). Overview of the CCP4 suite and current developments. Acta Crystallogr. Sect. D Biol. Crystallogr. 67, 235–242.

Yadav, M.K., Gerdts, C.J., Sanishvili, R., Smith, W.W., Roach, L.S., Ismagilov, R.F., Kuhn, P., and Stevens, R.C. (2005). In situ data collection and structure refinement from microcapillary protein crystallization. J. Appl. Crystallogr. 38, 900–905.

Zeldin, O.B., Gerstel, M., and Garman, E.F. (2013). RADDOSE-3D: time- and space-resolved modelling of dose in macromolecular crystallography. J. Appl. Crystallogr. 46, 1225–1230.

Zielinksi, K.A., Prester, A., Andaleeb, H., Bui, S., Yefanov, O., Catapano, L., Henkel, A., Wiedorn, M.O., Lorbeer, O., Crosas, E., et al. (2022). Rapid and efficient room temperature serial synchrotron crystallography using the CFEL TapeDrive. IUCrJ 9, 778–791.

